# Accessing Enzyme Kinetic Data and Prediction Methods at Scale

**DOI:** 10.64898/2026.09.25.751968

**Authors:** Saleh Alwer, Hugues Escoffier, Karim Taha, Veda Boorla, Han Yu, Somtirtha Santra, Zechen Wang, Chidi Egwu, Abraham Osinuga, Supantha Dey, Vaishnavey Srinivasan Raghunath, Farid Zare, Jack McGoldrick, Jan-Niklas Weder, Eduard Kerkhoven, Xiaozhou Luo, Costas D. Maranas, Liangzhen Zheng, Ulrike Wittig, Ratul Chowdhury, Rajib Saha, Nadine Töpfer, Thomas Sauter, Ronan M. T. Fleming

## Abstract

Enzyme kinetic parameters inform metabolic models, yet experimental measurements are sparse. A growing body of work predicts them from protein and substrate features, but software fragmentation hinders adoption, so downstream tools lock into the most accessible method. We present OpenKinetics Predictor (<u>predictor.openkinetics.org</u>), an open-source platform integrating thirteen kinetic parameter prediction methods accessible via one interface. The platform optionally reports similarity between query proteins and the training data of each method to provide context. A shared framework for featurisation and prediction keeps it extensible, and independent parties, including original authors, contributed many methods. A paired data portal (<u>data.openkinetics.org</u>) exposes a curated kinetic dataset with precomputed embeddings, predicted binding sites, and standardised splits. Both offer a web interface and an API, and the GECKO modelling toolbox already calls the predictor API. As a case study, we predict across an *E. coli* model and find inter-predictor agreement varies with metabolic context and data availability.

## INTRODUCTION

Kinetic parameters constrain the rate of enzyme catalysed reactions. The turnover number (*k*_cat_) bounds the maximum rate per enzyme molecule, and the Michaelis constant *(K*_M_) gives the substrate concentration at half-maximal rate. Genome-scale metabolic models that incorporate these parameters predict growth, reaction fluxes, and protein allocation more accurately than models built on reaction stoichiometry alone^1–3^. Direct measurement is slow and costly, so experimental coverage reaches a small fraction of known enzymes. Curated repositories such as BRENDA^4^ and SABIO-RK^5^ hold values for tens of thousands of enzyme-catalysed reactions, but that is a small subset of the millions of enzyme coding sequences.

A growing set of computational methods predict kinetic parameters from protein sequences and substrate structures, drawing on protein language model embeddings and molecular fingerprints^3,6–18^. These methods extend kinetic coverage to enzymes that have not been experimentally characterised. Adoption lags capability because each method ships with its own input format, software dependencies, output structure, and execution assumptions. Installing and running several methods for comparison demands sustained engineering effort. Downstream modelling and engineering workflows therefore adopt a single predictor, often the one most readily available rather than the one best suited to the proteins under study, and rarely switch to methods published later. A downstream tool locked to an older predictor underperforms relative to what a newer, more accurate one allows^6^. This lock-in also hides a question central to model building: do different predictors agree, and where they disagree, does the disagreement follow a structure a modeller should account for? Such disagreement is one facet of the broader, often unquantified uncertainty limiting confidence in these models^19^.

We present OpenKinetics Predictor, an open-source platform integrating thirteen published prediction methods accessible via one interface. The platform standardises input and output, runs each method in an isolated environment, and exposes both a web interface and an application programming interface (API). On request, queries are matched against experimental values in BRENDA^4^, SABIO-RK^5^, and UniProt^20^, and an experimental value can optionally overwrite the corresponding prediction. The platform also reports the sequence similarity between each query protein and the training data of each method, as sequence similarity has been shown to correlate with prediction accuracy. The enzyme-constrained modelling toolbox GECKO^2^ originally used DLKcat^15^ for *k*_cat_ predictions and now calls the OpenKinetics Predictor API, so modellers can select the predictor. A shared framework for featurisation and prediction keeps the platform extensible, and independent groups, including several original method authors, contributed many of the integrated methods. Protein language model embeddings dominate the runtime of most methods. The platform caches these embeddings to speed up repeated queries and computes them on a remote GPU to speed up new ones. We pair the predictor with data.openkinetics.org, an open portal that exposes a curated kinetic dataset alongside precomputed protein embeddings, predicted binding sites, and standardised dataset splits for training and evaluation.

To examine predictor agreement at scale, we predict *k*_cat_ for every enzyme-reaction pair in the iML1515^21^ model of *Escherichia coli* using eight methods, and we measure their agreement. Predictions diverge widely, and the divergence is not random. Agreement rises with the availability of nearby experimental data and varies sharply with metabolic context, weakest in transport and membrane-associated reactions. These results mark where predictor choice carries the greatest consequence for systems-level modelling.

## RESULTS

### Platform architecture and unified interface

OpenKinetics Predictor exposes one interface to thirteen prediction methods (Figure 1, Table 1). A user supplies protein sequences and substrate structures in a single standardised CSV or JSON file, selects the prediction targets (*k*_cat_, *K*_M_, *k*_cat_/*K*_M_) and chooses a method for each target. Two options control the output: appending sequence-similarity columns and prioritising measured values over predictions. The platform validates inputs, parses SMILES and InChI substrate strings, enforces sequence-length limits, and checks method-specific constraints before dispatch (Figure 1A).

**Figure 1.**
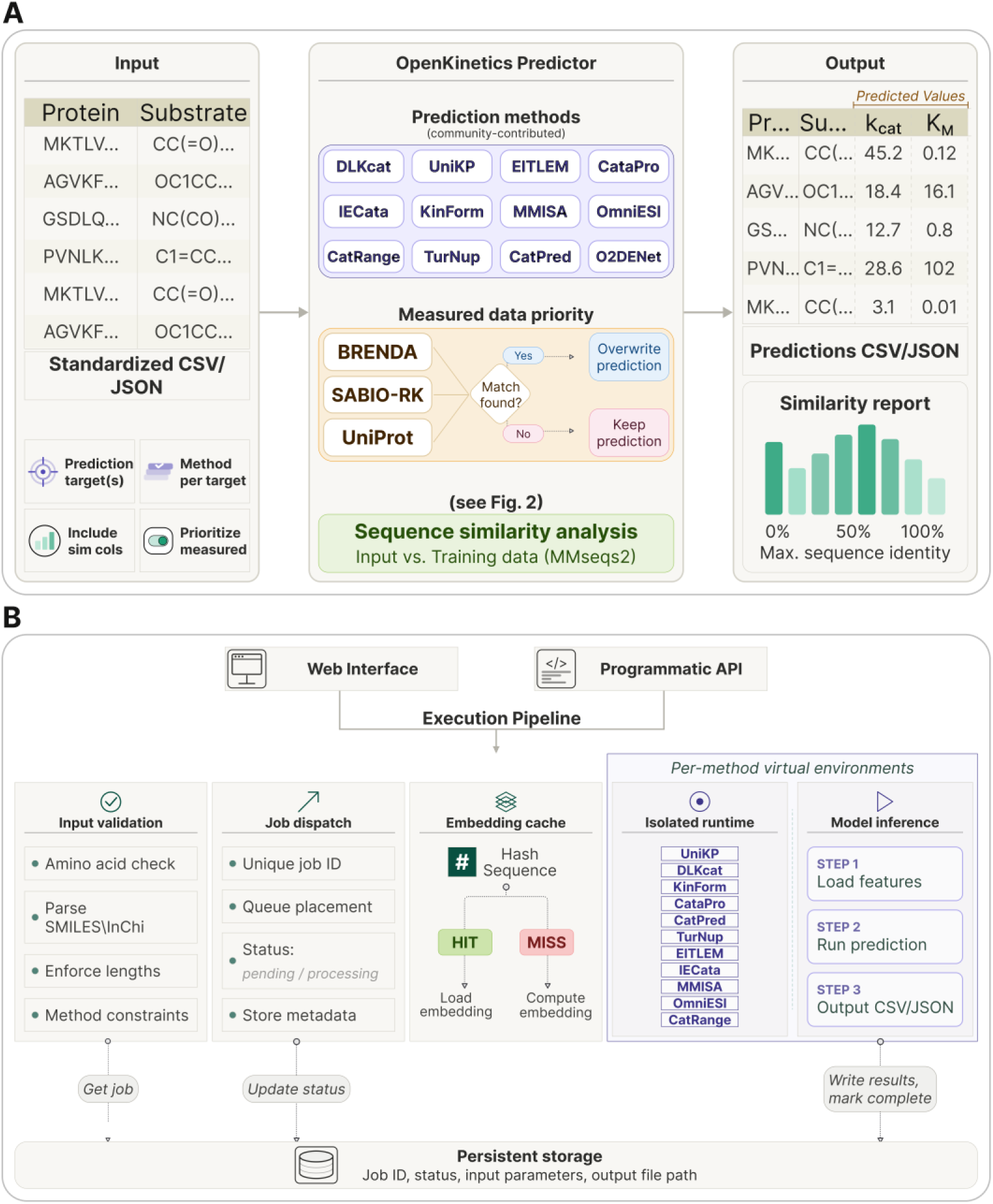
OpenKinetics Predictor provides unified access to enzyme kinetic parameter prediction methods. The platform provides unified access to multiple enzyme kinetic parameter prediction methods through one standardised interface, handling input validation, method execution, and output generation. (A) Conceptual workflow. Users supply protein sequences and substrate structures as a standardised CSV or JSON file, together with the prediction targets (*k*_cat_, *K*_M_, *k*_cat_/*K*_M_) the method selected per target, and options to include similarity columns and prioritise measured values. The platform routes inputs to the chosen method(s). When measured values are requested, queries are matched against experimental databases (BRENDA, SABIO-RK, UniProt) and, where a match is found, the measured value overwrites the prediction. An optional sequence similarity analysis compares input proteins against each method’s training data using MMseqs2 (see Figure 2). The output is returned as a predictions table (CSV or JSON) with kinetic parameter values, accompanied by similarity results if requested. (B) System architecture. Jobs submitted through the web interface or programmatic API enter an execution pipeline, which validates inputs, dispatches each job with a unique identifier, and records metadata in persistent storage. Protein embeddings are retrieved from a cache when available (HIT) or computed and stored when absent (MISS). Each prediction method runs in its own isolated virtual environment, and completed predictions are written back to persistent storage.

**Table 1.** Overview of the prediction methods currently available. Supported kinetic parameters are marked with a checkmark. Benchmark runtimes were measured on 1,000 reactions involving 100 unique protein sequences. Runtimes are reported with protein language model embeddings computed on GPU or CPU without caching, and with pre-computed embeddings available in the cache. NA indicates the configuration does not apply to the method.

| Method | Publication | Parameter |  |  | Not cached |  | Cached |
| --- | --- | --- | --- | --- | --- | --- | --- |
| | | $k_{\text{cat}}$ | $K_M$ | $k_{\text{cat}}/K_M$ | GPU | CPU | |
| DLKcat | Li et al., Nat. Catal. (2022)<sup>15</sup> | ✓ |  |  | NA | 32 s | NA |
| MMISA-KM | Song & Wang, IEEE DDCLS (2025)<sup>12</sup> |  | ✓ |  | NA | 4 min 22 s | NA |
| CatPred | Boorla & Maranas, Nat. Commun. (2025)<sup>17</sup> | ✓ | ✓ | ✓ | 5 min 54 s | 14 min 0 s | 23 s |
| CatRange (orig. RealKcat) | Sajeevan et al., PNAS Nexus (2026)<sup>18</sup> | ✓ |  |  | 1 min 38 s | 14 min 43 s | 1 min 9 s |
| EITLEM-Kinetics | Shen et al., Chem Catal. (2024)<sup>14</sup> | ✓ | ✓ |  | 7 min 43 s | 18 min 13 s | NA |
| OmniESI | Nie et al., arXiv (2025)<sup>11</sup> | ✓ | ✓ |  | 7 min 34 s | 18 min 41 s | NA |
| OmniESI + O2DENet | Wu et al., J. Chem. Inf. Model. (2026)<sup>29</sup> | ✓ | ✓ |  | 7 min 34 s | 18 min 41 s | NA |
| TurNuP | Kroll et al., Nat. Commun. (2023)<sup>8</sup> | ✓ |  |  | 3 min 53 s | 19 min 36 s | 2 min 12 s |
| CataPro | Wang et al., Nat. Commun. (2025)<sup>13</sup> | ✓ | ✓ | ✓ | 1 min 37 s | 25 min 9 s | 41 s |
| UniKP | Yu et al., Nat. Commun. (2023)<sup>7</sup> | ✓ | ✓ |  | 1 min 28 s | 33 min 46 s | 58 s |
| IECata | Wang et al., Brief. Bioinform. (2025)<sup>10</sup> | ✓ |  |  | 11 min 11 s | 34 min 41 s | NA |
| | | $k_{\text{cat}}$ | $K_{\text{M}}$ | $k_{\text{cat}}/K_{\text{M}}$ | GPU | CPU | |
| KinForm-L | Alwer & Fleming,<br>npj Syst. Biol. Appl.<br>(2026)<sup>6</sup> | ✓ |  |  | 3 min 49<br>s | 54 min<br>38 s | 37 s |
| KinForm-H | Alwer & Fleming,<br>npj Syst. Biol. Appl.<br>(2026)<sup>6</sup> | ✓ | ✓ |  | 3 min 42<br>s | 56 min<br>10 s | 36 s |

Each job receives a unique identifier and enters an execution pipeline (Figure 1B). Protein language model embeddings dominate runtime for most methods, so the platform caches them keyed on a hash of the sequence. A cache hit reuses a stored embedding, and a miss computes and stores a new one. Embedding computation can run on a remote GPU server dedicated to this step. Each method then runs in its own isolated virtual environment, which removes the dependency conflicts otherwise blocking side-by-side use. Completed predictions and job metadata are written to persistent storage and returned through the web interface or API.

The integrated methods differ in which targets they predict and in their speed (Table 1). Runtime spans two orders of magnitude across methods. Caching provides large speedups on repeated queries, in several cases from tens of minutes to under one minute. Computing embeddings on a GPU rather than a CPU speeds up new queries. When a user requests measured values, the platform queries BRENDA, SABIO-RK, and UniProt, and on a match the experimental value overwrites the prediction (Figure 1A).

### Sequence similarity contextualises prediction reliability

Sequence similarity to training data provides context for prediction reliability, as prediction accuracy has been shown to decline as query proteins become more distant from the data used for training^6,8,22^. OpenKinetics Predictor therefore reports the similarity between each query protein and the training data available for each method (Figure 2).

**Figure 2.**
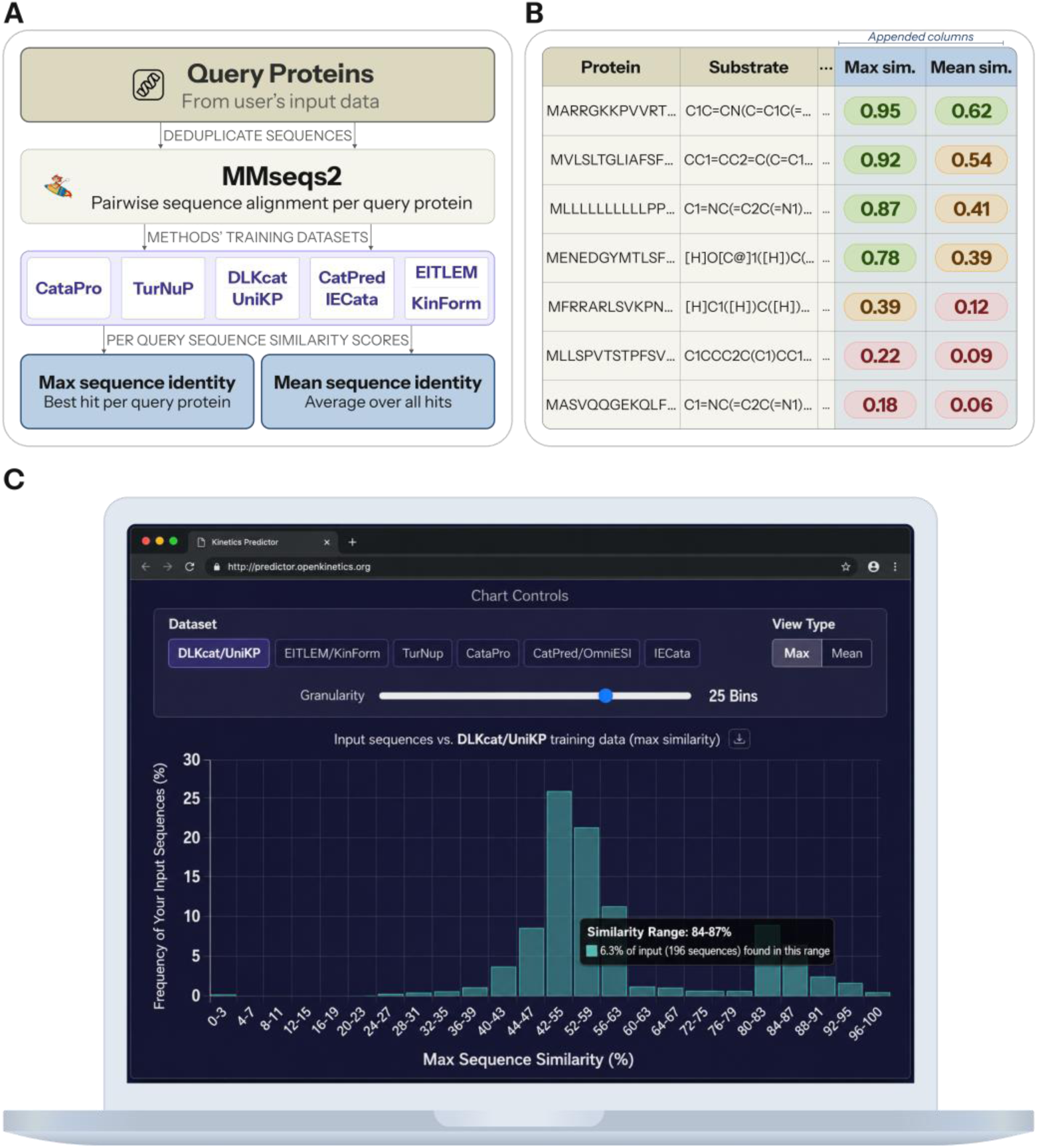
Sequence similarity to training data contextualises prediction reliability. Query protein sequences are deduplicated and aligned with MMseqs2 against the training datasets of the supported prediction methods, quantifying how similar user inputs are to each method’s training data. (A) For each query protein, the pipeline computes two summary measures for each training dataset: maximum sequence identity (the best hit) and mean sequence identity (the average across all hits). (B) These similarity scores are appended as additional columns to the output table, so each prediction row carries its corresponding maximum and mean similarity values. (C) The web interface provides an interactive similarity histogram summarising the distribution of similarity scores across all input proteins for a selected training dataset. Controls switch between datasets, toggle between maximum and mean similarity, and adjust the histogram binning.

Each prediction is accompanied by maximum and mean sequence identity to the relevant training set (Figure 2A,B), allowing users to interpret predictions in the context of how closely the query resembles proteins represented during training. The web interface also summarises these similarities across all input proteins as an interactive histogram (Figure 2C). Query sets concentrated at low sequence identity indicate greater extrapolation beyond the training data, while higher identity indicates closer representation within the training set.

### Predictors disagree substantially across the iML1515 *E. coli* model

To test how predictor choice affects a genome-scale model, we predicted *k*_cat_ for every enzyme-reaction pair in iML1515^21^ using eight method configurations (Figure 3). We compared predictions on a log_10_ scale. Pairwise Pearson correlation ranged from *r* = 0.08 to *r* = 0.67 (Figure 3, upper triangle). The strongest agreement was between the two KinForm configurations (*r* = 0.67), which share architecture and training data. The weakest was between DLKcat and TurNuP (*r* = 0.08). The marginal distributions also differed in centre and spread (Figure 3, diagonal), and the scatter plots show wide deviation from the line of perfect agreement (Figure 3, lower triangle). A correlation of 0.67 between the most similar pair leaves substantial unexplained variance, and most pairs agreed far less. The kinetic parameters assigned to a model therefore depend heavily on which predictor is used.

**Figure 3.**
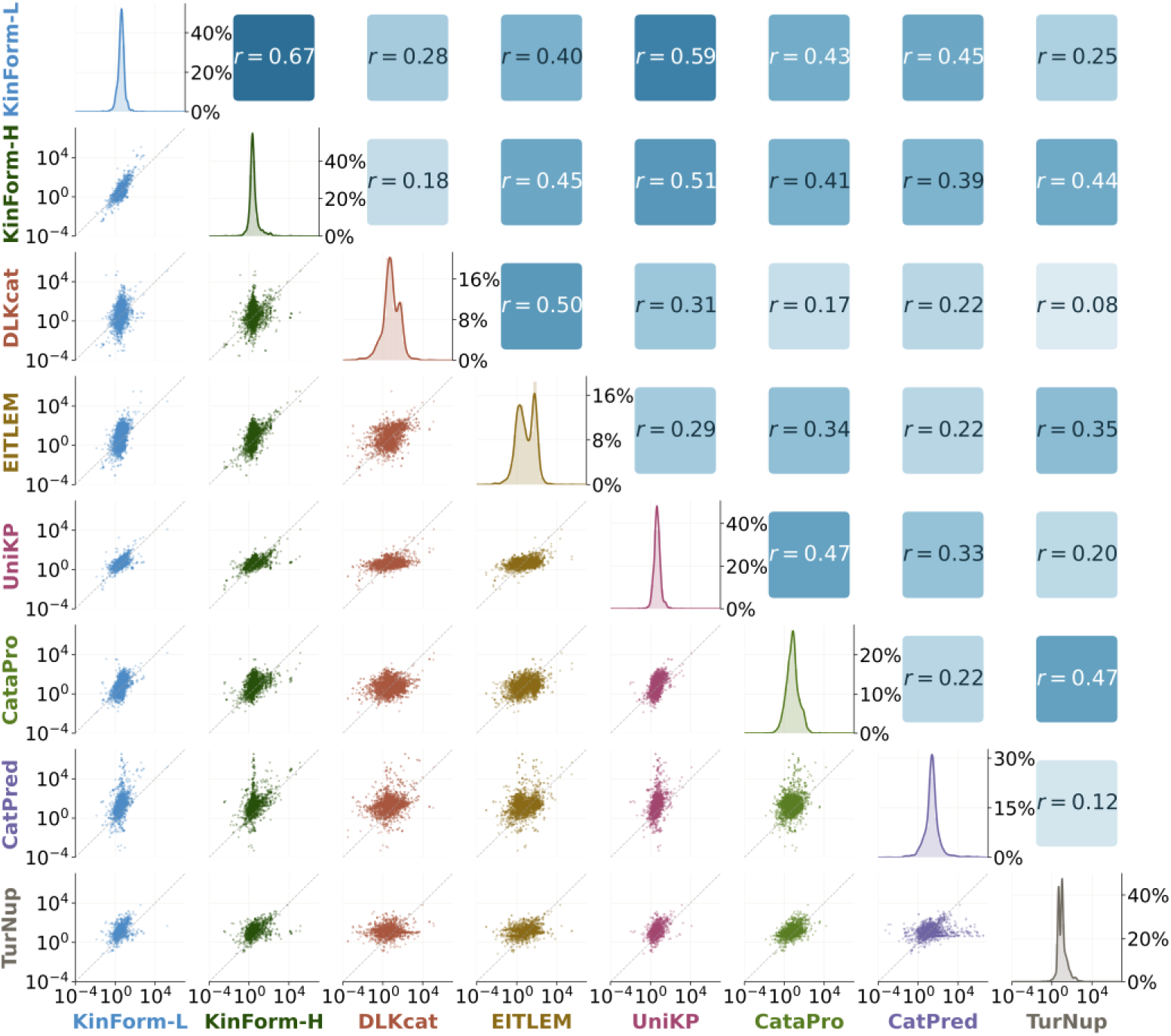
Predicted *k*_cat_ values disagree substantially across methods in the iML1515 *E. coli* model. Predicted log_10_(*k*_cat_) values from eight method configurations compared for all enzyme-reaction pairs in the iML1515 metabolic model. (Diagonal) Distribution of each method’s predictions across the model. (Lower triangle) Pairwise scatter plots of predicted log_10_(*k*_cat_) values. The dashed grey line marks perfect agreement. (Upper triangle) Pearson correlation coefficient (*r*) for each pair, computed on log-transformed predictions, with cell shading scaled to *r*. Correlations span a wide range (*r* = 0.08 to 0.67), showing methods often disagree substantially on the same enzyme-reaction pairs.

### Agreement is structured by data availability and metabolic context

We grouped enzyme-reaction pairs by the WILDkCAT^24^ penalty score, which reflects how closely each pair matches an entry in BRENDA or SABIO-RK (Figure 4A). Mean correlation with the other methods was highest at low penalty scores and declined as experimental support weakened. Predictor choice therefore matters most in the pathways where experimental guidance is thinnest. Agreement also tracked metabolic context (Figure 4B). Within amino acid metabolism and within energy production and conversion, correlations were strong across most method pairs. Within transport and outer or inner membrane reactions, correlations were weak and, in several pairs, negative.

**Figure 4.**
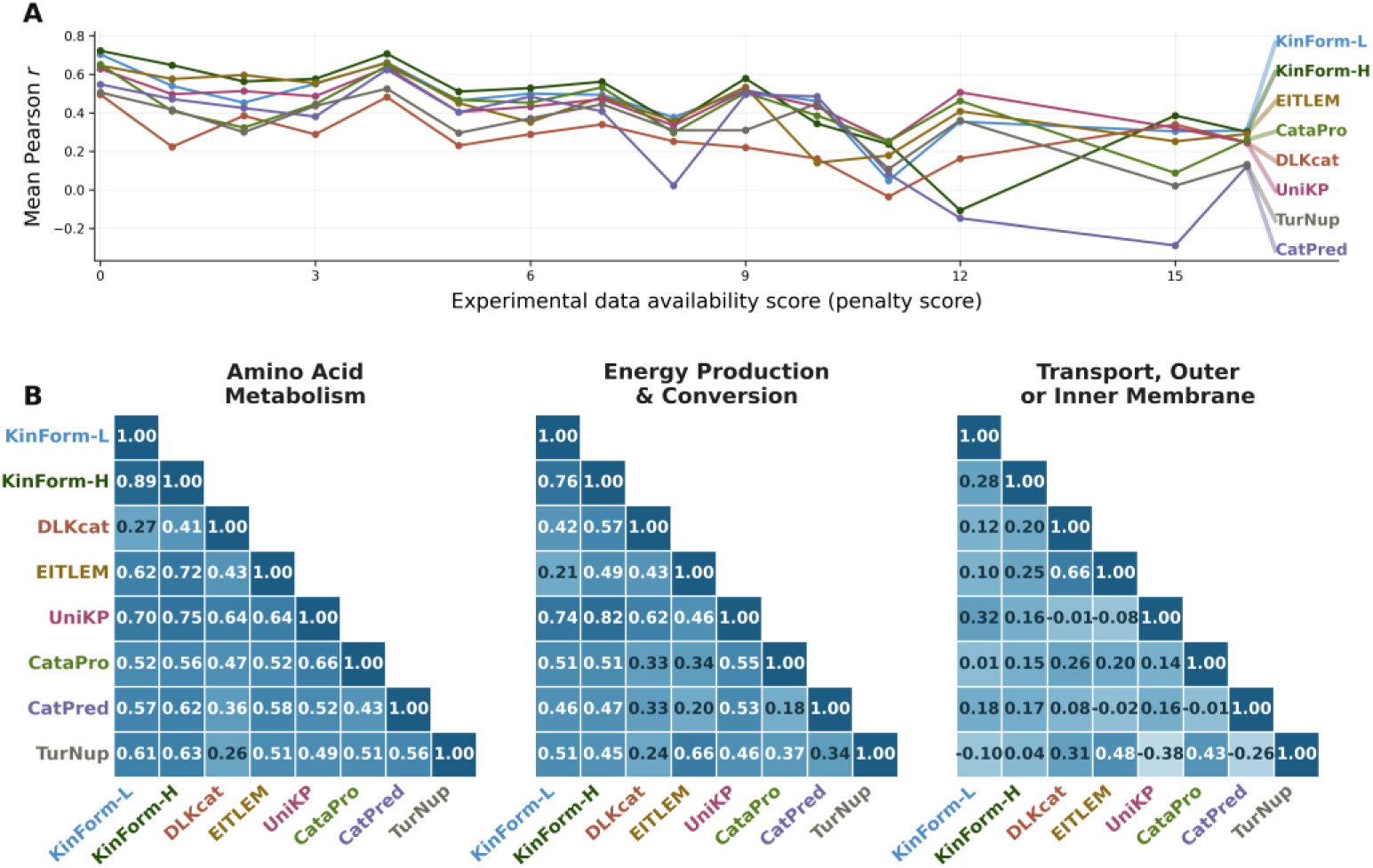
Inter-method agreement is structured by experimental data availability and metabolic context in *E. coli*. Pairwise Pearson correlation between *k*_cat_ predictions across eight method configurations, computed over enzyme-reaction pairs in iML1515. (A) Mean correlation of each method with all others, grouped by the best-match penalty score of each enzyme-reaction pair in the experimental databases (BRENDA and SABIO-RK). The penalty score^24^ reflects experimental data availability: 0 marks an exact match in the database, and higher scores mark progressively poorer matches, up to 16 where no experimental value exists. Agreement is highest at low penalty scores and declines as experimental support weakens. (B) Pairwise correlation matrices within three metabolic pathway categories. Cell shading is scaled to correlation. Agreement is strong in amino acid metabolism and in energy production and conversion, and weak or negative in transport, outer or inner membrane.

### Agreement with in-distribution measured values

For the 142 enzyme-reaction pairs in iML1515 with an experimental match, we compared predictions against measured *k*_cat_ values (Figure 5), using RMSE, MAE, and Pearson correlation on log_10_-transformed values. RMSE ranged from 0.93 to 1.40 and r from 0.44 to 0.71 across the eight configurations. This comparison must not be read as a general accuracy ranking. It measures in-distribution performance only. The experimental values come from the same database family used to train most methods, so methods designed for out-of-distribution prediction appear less favourable here. The comparison describes fit to familiar data, and a method placed low here can still generalise better to unfamiliar enzymes.

**Figure 5.**
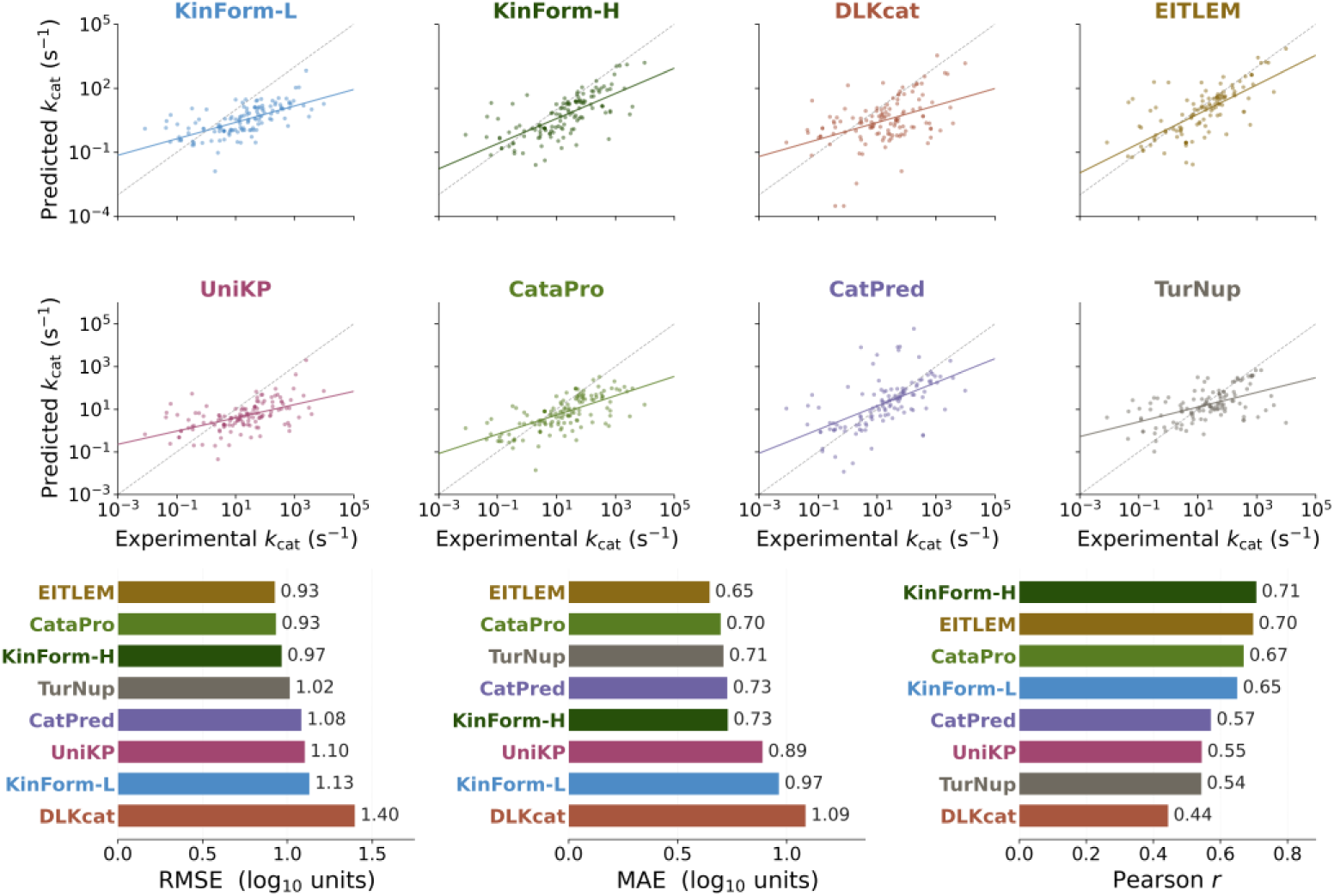
Predicted versus in-distribution measured *k*_cat_ values in *E. coli*. Predictions from eight method configurations compared against experimental *k*_cat_ measurements for enzyme-reaction pairs in iML1515 with a database match. (Top) Predicted versus experimental log_10_(*k*_cat_) for each method. The solid-coloured line is the linear regression fit and the dashed grey line marks perfect agreement. (Bottom) Method ranking on three metrics: root mean square error (RMSE), mean absolute error (MAE), and Pearson correlation coefficient (*r*). Lower RMSE and MAE and higher *r* indicate greater accuracy. All are calculated on log_10_-transformed values. These results reflect in-distribution performance, not generalisation ability. The experimental *k*_cat_ values come from the same database family used for training. Models designed for out-of-distribution prediction therefore appear less favourable here.

### An open data portal for kinetic data and features

Alongside the predictor, we host data.openkinetics.org, an open portal for curated enzyme kinetic data and precomputed features. The portal exposes CatLog, a manually curated, agentically corroborated with human-in-the-loop, dataset of 50,312 kinetic parameters compiled by the CatRange (orig. RealKcat) team^18^, to be published separately. For every protein sequence in a release, the portal provides precomputed language model embeddings from ESMC^25^, ESM2^26^, and ProtT5^27^, and predicted binding sites from Pseq2Sites^28^. It also provides train, test, and validation splits under three schemes: random, sequence-exclusive, and substrate-exclusive, where an exclusive split places no identical sequence, or no identical substrate, in more than one partition. The portal is versioned, and each release ships its own data, features, and splits. These downloads remove repeated preprocessing and give methods a common basis for training and evaluation.

## DISCUSSION

We developed OpenKinetics Predictor to make kinetic parameter prediction methods accessible through one interface. Applying these methods to a genome-scale model showed that predicted values differ substantially between predictors, even though the methods train on overlapping data, and that this divergence concentrates in specific parts of metabolism. Prior work shows that the choice of predictor propagates into downstream modelling tasks such as phenotype prediction^6,8,15^. The choice of predictor is therefore a modelling decision with measurable consequences.

Inter-method agreement offers a way to assess predictions in the absence of measured values. Where methods converge, the predicted parameter is stable with respect to the choice of method. Where methods diverge, the value assigned to a reaction depends largely on which method was selected. Agreement should not be interpreted as correctness. Most published methods are trained on overlapping subsets of BRENDA and SABIO-RK, so their errors are correlated and they can agree while remaining incorrect. Agreement provides a measure of how sensitive a parameterisation is to the choice of prediction method, without requiring experimental data.

Interpreting any comparison of methods needs a measure of how far a query sits from a method’s training data. Reporting the sequence identity between query proteins and each method’s training data provides this context, and we make the measure available for any input dataset. This is analogous to the applicability domain used in quantitative structure-activity relationship modelling and to per-residue confidence scores in protein structure prediction.

For enzyme-constrained metabolic modelling, the results identify that transport and membrane-associated reactions show low inter-method agreement and are underrepresented in kinetic databases, and the proteins that catalyse them differ structurally from the soluble enzymes that dominate training data. Models of phenotypes that depend on transport capacity therefore rest on weaker kinetic support than models of central carbon metabolism. The same analysis identifies which enzymes would most improve model reliability if measured, which provides a basis for prioritising experimental effort. Earlier work found that the choice of turnover number predictor shifts the output of downstream enzyme-constrained modelling tasks, although the differences were modest and the comparison covered a small number of methods^6,8,15^. Together, these results suggest that predictor choice may influence downstream model behaviour. In this work, we did not quantify how disagreement between predictors propagates into predicted fluxes, growth rates, or protein allocation, and this is a direct extension of the present work.

The platform supports analyses that span several methods at once. Predictions can be combined across methods, measured values can be substituted where experiments exist, and different methods can be applied to different protein classes within one model. GECKO^2^, an enzyme-constrained metabolic modelling toolbox, can now call the OpenKinetics Predictor API. A modeller selects a predictor instead of the hard-coded DLKcat default, and GECKO records the source of each prediction for downstream tuning. The platform also lowers the cost of evaluating new methods and makes them more accessible, since integration requires only the two stages of the featurisation-prediction abstraction. Several methods were integrated by collaborators across institutions, demonstrating that the framework supports distributed contributions. The open-source release includes the container definitions for every component, so any group can host the full platform on their own server if needed.

### Limitations of the study

The case study covers a single organism. We applied the methods to iML1515, a high-quality genome-scale model of *Escherichia coli*, the best characterised bacterial species^21^. Inter-method agreement and the accuracy of individual methods may differ in organisms with less experimental coverage, and the pathway-level pattern reported here should not be assumed to hold in other species.

The case study used eight of the thirteen integrated method configurations, since the remainder do not predict *k*_cat_ or were unavailable at the time of the analysis. The agreement analysis therefore describes these eight methods and not the full set available through the platform.

The accuracy comparison is in-distribution, because the reference values come from the same databases that trained the methods, so the comparison is not a general measure of accuracy. A controlled evaluation would retrain each method on one training set and test it on one held-out set, separating the method from its training data and enabling a like-for-like comparison. Building and running such an evaluation is beyond the scope of this paper. The sequence-exclusive and substrate-exclusive splits released through data.openkinetics.org provide a starting point for such an evaluation.

Inter-method agreement measures consistency, not accuracy. We quantified it as the Pearson correlation of log-transformed predictions, which captures co-variation across reactions but not systematic offsets in absolute magnitude. Two methods can correlate strongly while differing in scale, and methods trained on overlapping data can agree while all being incorrect.

The runtime benchmark reflects each method’s original implementation. Every method was integrated from its published repository without rewriting its internals, and these repositories differ in how well they parallelise work and exploit GPU acceleration. Reported runtimes therefore measure the code as released, not the method’s inherent speed. Optimising each implementation to a common standard would give a fairer comparison.

We did not validate predictions experimentally, and we did not test how the choice of method affects downstream model behaviour. Whether disagreement between predictors changes predicted fluxes, growth rates, or protein allocation in an enzyme-constrained model remains to be determined.

The platform runs as a hosted web service and API. We do not yet provide an installable package for local use inside an analysis script, and this is a direction for future work.

## RESOURCE AVAILABILITY

### Lead contact

Requests for further information and resources should be directed to and will be fulfilled by the lead contact, Ronan M. T. Fleming.

### Materials availability

This study did not generate new unique reagents.

### Data and code availability

- Data: the iML1515 *k*_cat_ predictions and case-study outputs have been deposited at Zenodo. The DOI is listed in the key resources table.
- Code: the source code is hosted on GitHub, with the release for this paper archived at Zenodo. The DOI is listed in the key resources table.

## ACKNOWLEDGMENTS

This work was supported by the European Union’s Horizon Europe programme (grant 101080997), the Swiss State Secretariat for Education, Research and Innovation, SERI (grant 23.00232), UK Research and Innovation, UKRI (grants 10083717 and 10080153), the Luxembourg National Research Fund, FNR (grant PRIDE21/16763386/CANBIO2), the Novo Nordisk Foundation (grants NNF10CC1016517 & NNF20CC0035580), the Knut and Alice Wallenberg Foundation, the European Union’s Horizon 2020 programme (grants 686070 and 814650), China’s National Key R&D Programme (grant 2025YFA0922700), and the German Research Foundation (DFG) under Germany’s Excellence Strategy (EXC-2048/1, project ID 390686111) and SFB 1644/1 (project no. 512328399).

## AUTHOR CONTRIBUTIONS

Conceptualization, S.A., E.K., and R.M.T.F.; Methodology, S.A. and H.E.; Software, S.A., K.T., V.B., H.Y., S.S., Z.W., C.E., J.M., A.O., and V.S.R.; Formal Analysis, S.A. and H.E.; Investigation, H.E., F.Z., and E.K.; Resources, E.K., U.W., R.C., and R.S.; Data Curation, S.A., H.E., A.O., V.S.R., R.C., S.D., and R.S.; Writing – Original Draft, S.A.; Writing – Review & Editing, all authors; Visualization, S.A., H.E., A.O., V.S.R., R.C., and R.S.; Supervision, J.-N.W., N.T., X.L., C.D.M., L.Z., T.S., R.C., R.S., and R.M.T.F.; Project Administration, S.A. and R.M.T.F.; Funding Acquisition, E.K., X.L., C.D.M., L.Z., U.W., R.C., R.S., N.T., T.S., and R.M.T.F.

## DECLARATION OF GENERATIVE AI AND AI-ASSISTED TECHNOLOGIES IN THE WRITING PROCESS

During the preparation of this work, we used Claude to check sentence structure and grammar. We also used Claude to review the codebase against the Methods section and identify any implementation details that had been omitted. After using this tool, we reviewed and edited the content as needed. We take full responsibility for the content of the publication.

## STAR★METHODS

### KEY RESOURCES TABLE

STAR★METHODS KEY RESOURCES TABLE
| REAGENT OR RESOURCE | SOURCE | IDENTIFIER |
| --- | --- | --- |
| ***Antibodies*** |  |  |
| N/A |  |  |
| ***Bacterial and virus strains*** |  |  |
| N/A |  |  |
| ***Biological samples*** |  |  |
| N/A |  |  |
| ***Chemicals, peptides, and recombinant proteins*** |  |  |
| N/A |  |  |
| ***Critical commercial assays*** |  |  |
| N/A |  |  |
| ***Deposited data*** |  |  |
| iML1515 $k_{cat}$ predictions and case-study outputs | This paper | Zenodo: 10.5281/zenodo.21370513 |
| Original code, OpenKinetics Predictor | This paper | Zenodo: 10.5281/zenodo.21361461 |
| BRENDA enzyme database | Chang et al., 2021<sup>4</sup> | www.brenda-enzymes.org;<br>RRID:SCR_002997 |
| SABIO-RK | Wittig et al., 2018<sup>5</sup> | sabiork.h-its.org; |
| UniProt | UniProt Consortium, 2025<sup>20</sup> | www.uniprot.org; RRID:SCR_002380 |
| iML1515 genome-scale model of *E. coli* | Monk et al., 2017<sup>21</sup> | BiGG Models:<br>bigg.ucsd.edu/models/iML1515 |
| CatLog dataset | CatRange (orig. RealKcat) team<sup>18</sup>, to be published | chowdhurylab.github.io/tools/catlog-latest.html;<br>data.openkinetics.org; |
| **Experimental models: Cell lines** |  |  |
| N/A |  |  |
| **Experimental models: Organisms/strains** |  |  |
| N/A |  |  |
| **Oligonucleotides** |  |  |
| N/A |  |  |
| ***Recombinant DNA*** |  |  |
| N/A |  |  |
| ***Software and algorithms*** |  |  |
| OpenKinetics Predictor (web interface and API) | This paper | predictor.openkinetics.org;<br>github.com/openKinetics/webKinPred;<br>Zenodo:<br>doi.org/10.5281/zenodo.21361461 |
| Data Portal | This paper | data.openkinetics.org;<br>github.com/openKinetics/openkinetics-data; |
| DLKcat | Li et al., 2022<sup>15</sup> | github.com/SysBioChalmers/DLKcat;<br>v1.0.0 |
| UniKP | Yu et al., 2023<sup>7</sup> | github.com/Luo-SynBioLab/UniKP; v1 |
| TurNuP | Kroll et al., 2023<sup>8</sup> | github.com/AlexanderKroll/Kcat_prediction;<br>Initial release for TurNuP paper |
| CatPred | Boorla and Maranas, 2025<sup>17</sup> | github.com/maranasgroup/CatPred;<br>b314a28 |
| CataPro | Wang et al., 2025<sup>13</sup> | github.com/zchwang/CataPro/; cc89b2c |
| EITLEM-Kinetics | Shen et al., 2024<sup>14</sup> | github.com/XvesS/EITLEM-Kinetics;<br>7bb9d66 |
| KinForm-L and KinForm-H | Alwer and Fleming, 2026<sup>6</sup> | github.com/Digital-Metabolic-Twin-Centre/KinForm; f7a70eb |
| MMISA-KM | Song and Wang, 2025<sup>12</sup> | github.com/kaiwang-group/MMISA-KM;<br>f1783b6 |
| CatRange<br>(orig.<br>RealKcat) | Sajeevan et al., 2026<sup>18</sup> | github.com/ssbio/CatRange;<br>9e4336b |
| OmniESI | Nie et al., 2025<sup>11</sup> | github.com/Hong-yu-Zhang/OmniESI;<br>de7d027 |
| IECata | Wang et al., 2025<sup>10</sup> | github.com/zhaoyanpeng208/IECata;<br>cce38a4 |
| O2DENet | Wu et al., 2026<sup>29</sup> | github.com/blackjack534/O2DENet;<br>cb02c38 |
| MMseqs2 | Steinegger and Söding,<br>2017<sup>23</sup> | RRID:SCR_022962; release 13.45111 |
| ESM | Lin et al., 2023<sup>26</sup> | esm2_t33_650M_UR50D |
| ProtT5 | Elnaggar et al., 2022<sup>27</sup> | prot_t5_xl_uniref50 |
| Django | Django Software<br>Foundation<sup>30</sup> | djangoproject.com; version 5.1.15 |
| Celery | Celery project<sup>31</sup> | docs.celeryq.dev; version 5.6.3 |
| Redis | Redis Ltd.<sup>32</sup> | redis.io; version 7.4.8 |
| SQLite | Hipp, D.R., and Team<sup>33</sup> | sqlite.org; version 3.46.1 |
| Python | Python Software<br>Foundation<sup>34</sup> | python.org; version 3.12.13 |
| COBRApy | Ebrahim et al., 2013<sup>35</sup> | RRID:SCR_012096; version 0.29.0 |
| **Other** |  |  |
| N/A |  |  |

### METHOD DETAILS

#### Platform architecture

OpenKinetics Predictor runs a client-server-worker architecture with four components. A Python Django^30,34^ backend receives each job from the web interface or the API, validates the input, and writes a job record with a unique identifier to a SQLite database^33^. The backend places the job on a Celery^31^ task queue with Redis^32^ as the message broker. A worker retrieves the job, loads the input, runs method-specific preprocessing, activates the virtual environment for the selected method, and runs the prediction method. Predictions are written to a CSV file, and the file path is recorded against the job identifier. Clients poll job status and download the output on completion.

#### Input validation and substrate parsing

Users supply protein sequences and substrate structures in one standardised CSV or JSON file. Substrates are given as SMILES or InChI strings. The backend checks that each protein sequence contains only valid amino acid characters, and that each molecular string is syntactically valid. It then enforces method-specific constraints, including the maximum protein sequence length accepted by the selected method.

#### Sequence length handling

Sequences longer than a method’s limit are skipped or truncated, at the user’s choice. Truncation keeps the first *n/2* and the last *n/2* residues, where *n* is the maximum length for the selected method. This rule follows the long-sequence protocol used by Pseq2Sites^28^.

#### Input formats and row expansion

Methods accept one of three input types: a single protein-substrate pair, a protein with a substrate list, or a full reaction with substrates and products. Each method declares its type, and the platform routes inputs accordingly. A user can apply a single-substrate method to a multi-substrate or full-reaction input. The platform then expands the row, predicts for each substrate separately, and reduces the results to one value per row, taking the maximum for *k*_cat_ and *k*_cat_*/K*_M_ and the minimum for *K*_M_. This follows the convention set by the DLKcat paper. The platform handles enzyme complexes the same way. All methods natively predict for monomers, so a multimer is split into its subunits, each is predicted separately, and the best (maximum for turnover number and minimum for Michaelis constant) prediction is kept.

#### Protein embedding cache

Most methods featurise proteins with a protein language model, either ESM^26^ or ProtT5^27^ depending on the method. These embeddings dominate runtime for most methods. Each sequence maps to a stable internal identifier through a key-value store, and embeddings are held in method-specific directories. A worker loads a stored embedding when one exists, and computes and stores a new embedding otherwise. Substrate features, either molecular fingerprints or learned embeddings, are computed on demand because their cost is low.

#### Remote embedding service

Embedding computation can run on a separate GPU service that the backend contacts over HTTP. The backend sends the sequences whose embeddings are missing, waits for the embedding computation to complete, and verifies the cached artefacts before prediction. If the service is unavailable or a computation is incomplete, the backend falls back to local computation.

#### Isolated environments and the featurisation-prediction abstraction

Each method runs in its own virtual environment, which removes dependency conflicts between method codebases. Every method follows a common two-stage workflow. Featurisation converts sequences and substrates into numerical inputs. Prediction maps those inputs to a kinetic parameter. A new method integrates by implementing these two stages, which keeps the platform extensible.

#### Measured value lookup

When a user requests measured values, the platform queries BRENDA^4^, SABIO-RK^5^, and UniProt^20^ for the query enzyme and substrate. On a match, the measured value overwrites the corresponding prediction in the output and writes the relevant metadata.

#### Sequence similarity analysis

Prediction accuracy correlates with the similarity between a query sequence and a method’s training data^6,8,22^. The platform aligns query sequences against each method’s training set with MMseqs2^23^. Query sequences are deduplicated first. For each query and training set, the pipeline reports the maximum sequence identity, taken from the best hit, and the mean sequence identity, averaged across hits. These values attach to each prediction row as added columns, and the web interface summarises them via an interactive histogram across all input proteins.

#### iML1515 case study

We applied the eight configurations that predict *k*_cat_: KinForm-L, KinForm-H, DLKcat, EITLEM, UniKP, CataPro, CatPred, and TurNuP. We predicted one *k*_cat_ value for every enzyme-reaction pair in the iML1515 genome-scale metabolic model of *Escherichia coli*^21^. We used WILDkCAT^24^ to prepare the iML1515 model for prediction and to retrieve the experimental *k*_cat_ values used as reference. WILDkCAT maps a constraint-based metabolic model to enzyme-reaction queries, resolving gene-protein-reaction associations to enzyme sequences and assigning a substrate per reaction, and it retrieves measured wild-type *k*_cat_ values for these reactions from BRENDA and SABIO-RK under specified pH and temperature ranges. For each enzyme-reaction pair it reports a penalty score for the quality of the database match, where 0 marks an exact match and 16 marks no experimental value. We ran WILDkCAT on iML1515 to obtain one prediction query per enzyme-reaction pair, submitted these queries to OpenKinetics Predictor through its API, and used the retrieved *k*_cat_ values and penalty scores as the experimental reference for the accuracy and agreement analyses. Full details of the WILDkCAT pipeline and the penalty score are given in Escoffier et al.^24^.

#### Prediction comparison

We compared predictions on a log_10_ scale. Figure 3 compares all eight configurations across every enzyme-reaction pair. Figure 5 restricts the comparison to pairs with a database match, for which measured *k*_cat_ values are available. Statistical definitions appear in the quantification and statistical analysis section.

#### Data portal

The platform is accompanied by data.openkinetics.org, an open portal that hosts curated kinetic data and precomputed features. The curated dataset, CatLog, is produced by the CatRange (orig. RealKcat) team. Starting from BRENDA and SABIO-RK, they re-curate entries against the primary literature to correct errors, combining a manual verification campaign with agentic AI curation. The portal exposes CatLog for browsing and download. CatLog will be described in a separate publication by the authors. For each release, we precompute protein language model embeddings with ESMC, ESM2, and ProtT5, and predicted binding sites with Pseq2Sites, for every protein sequence in the dataset. We also provide train, test, and validation splits under three schemes. Random splits partition entries at random. Sequence-exclusive splits place no identical protein sequence in more than one partition. Substrate-exclusive splits place no identical substrate in more than one partition. The portal is versioned, and each release provides matching data, features, and splits.

### STUDY DESIGN

This study is computational. It used no experimental models and no study participants.

### QUANTIFICATION AND STATISTICAL ANALYSIS

We computed all correlations as Pearson correlation coefficients (r) on log_10_-transformed values. We report prediction error as root mean square error (RMSE) and mean absolute error (MAE), both in log_10_ units.

Figure 3 reports pairwise r across the eight configurations over the enzyme-reaction pairs in iML1515 for which both configurations in a pair returned a prediction. The number of jointly predicted pairs varies by configuration pair, from 3,577 to 4,567. Figure 4A reports each configuration’s mean r with the other seven, grouped by penalty score. Figure 4B reports pairwise r within three metabolic categories. Figure 5 reports RMSE, MAE, and r against measured *k*_cat_ values for the subset of pairs with a database match (n = 142). Per-panel detail appears in each figure legend.

We ran the analysis in Python, using NumPy^36^ for the statistics and COBRApy^35^ for the iML1515 model.

